# Localizing hidden regularities with known temporal structure in the EEG evoked response data

**DOI:** 10.1101/093922

**Authors:** Aleksandra Kuznetsova, Elena Krugliakova, Alexei Ossadtchi

## Abstract

In this paper we describe a novel data driven spatial filtering technique that can be applied to the ERP analysis in order to find statistically significant hidden differential activations in the EEG data. The technique is based on the known morphological characteristics of the response. Underlying optimization problem is formulated as a generalized Rayleigh quotient maximization problem. We supply our technique with a relevant randomization-based statistical test to assess the significance of the discovered phenomenon. Furthermore, we describe an application of the proposed method to the EEG data acquired in the study devoted to the analysis of the auditory neuroplasticity. We show how the mismatch negativity component, a tiny and short-lasting negative response that hallmarks the novel stimuli activating primary error-detection mechanisms, can be detected after filtration.

## 1 Introduction

Superfine temporal resolution is the most significant advantage the electroencephalography (EEG) and magnetoencephalography (MEG) techniques offer to cognitive neuroscientists [1]. With these techniques and using the evoked-response methodology (ERP) it became possible to discover and then reliably detect the mismatch negativity phenomenon (MMN) — a tiny and short-lasting negative response that hallmarks the novel stimuli activating primary error-detection mechanisms. Since then a plethora of studies investigating the presence of the MMN response in various paradigms and involving various sensory modalities appeared. Recent neuroscience advances predict the presence of the MMN like response not only in the primary sensory brain regions but also in the structures responsible for executive control, decision making and value encoding such as medial prefrontal cortex(mPFC), posterior ad anterior cingulate cortex (PCC, ACC). Despite the averaging implied by ERP approach, the activity of these deeper located brain regions when recorded by the non-invasive EEG and MEG sensors tends to get obscured by that of more superficial structures impinging on the array of sensors.

Here we describe a novel data driven spatial filtering technique that can be applied to the ERP data in order to find statistically significant hidden differential activations not otherwise seen in the sensor data. Based on the expected morphological characteristics of the response it allows to find a spatial filter to single out the sought response from the influence of other sources. We supply our technique with a relevant randomization-based statistical test to assess the significance of thus discovered phenomenon.

## 2 Observed data model

EEG or MEG data recorded by a *K*-sensor array during the *i*-th repetition of a cognitive task can be written as the following linear combination

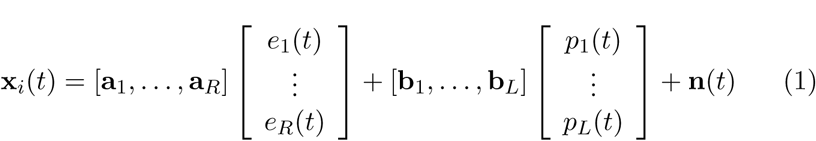

In other words, the recorded multichannel signal at each instance of time *t*, *x_i_*(*t*), is a noisy additive mixture of source topographies [**a**_1_,…, **a**_*R*_] weighted with the corresponding stimulus-locked activation timeseries [*e*_1_(*t*),…, *e*_*R*_(*t*)] and similarly represented task unrelated contribution from sources with topographies [**b**_1_,…, **b**_*L*_] scaled with [*p*_1_(*t*),…, *p_R_*(*t*)] time dependent activations. Topographies of task related sources form *K*-dimensional signal subspace and topographies of task unrelated sources form *L*-dimensional coherent interference subspace. ERP experiments are usually accompanied by a binary stimulus signal marking task onset. Usually the goal of data analysis is to identify task related signals and extract task-related signal component from these data. For completeness we should have included induced sources whose activation power is locked to the task-onset moment but the phase is random. However, since in the context of this paper we are interested in analysis of ERP we did not include the induced component in (1).

The averaging procedure emphasizes the phase-locked to the stimulus component of the response leaves us with the multichannel ERP data **X**(*t*)

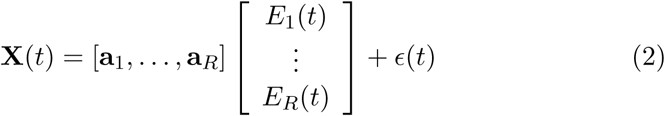

Source level ERPs *E_i_*(*t*) are stacked into the source timeseries vector that scales the corresponding source topographies **a**_*i*_. Source topography norms ∥**a**_*i*_∥ determine the signal-to-noise ratio specific to the *i*-th source. These norms vary due to location and orientation of the sources. The depth of a source appears to be the key factor here. For example, in the MEG data the field strength is inversely proportional to the cube of the source-sensor distance. In the EEG case the relation is complicated but results into significantly more pronounced potential spread of the deeper source as compared to the more superficial ones.

Taking into account the linearity of the above mixture, the activity of the *i*-th source can be estimated by means of spatial filtering that boils down to computing the linear combination *Ê_i_*(*t*) = **w**_*i*_**X**(*t*) of sensor signals with weights **w**_*i*_ determined by solving the inverse problem via one of the many available non-parametric, parametric techniques or by a data driven decomposition such as CSP [2] or ICA [6].

In the exploratory analysis the obtained source activation timeseries estimates *Ê_i_*(*t*) are then scrutinized for significance in order to draw conclusions related to the studied hypothesis. In the non-parametric scenario the entire inverse operator is obtained. The procedure of finding sources with significant activations requires a statistical test for multiple (~ 10^4^) hypotheses which results into a significant reduction of statistical power. When **w**_*i*_ is ob-tained by a spatial decomposition the multiple comparisons (~ 100) is still a problem and additionally, there is no guarantee the decomposition employed isolates the sought phenomenon into a single component. For instance, in [4] it has been demonstrated that the ICA tends to deliver a suboptimal performance in the task of finding the narrow-band sources when applied to a typical instantaneous mixture, i.e. (1). When solving the inverse problem with the parametric approach, the source of interest may be obscured by the activity of more superficial dipoles and therefore the dipole-fitting algorithm will fail to find this deeper source.

Response latency and peak duration are the most physiologically interpretable features of the ERPs and the corresponding source timeseries. Therefore, when searching for a neurophysiologically plausible solution it is often desirable to be able to supply this information to the algorithm. In what follows we describe a novel Rayleigh-quotient based method that allows to find weak sources whose activity possesses the desired morphological features.

## 3 Method

When performing comparative analysis of the responses in two conditions we usually have some expectation regarding the timing of the expected difference. For example, when studying classical MMN responses we expect the difference to occur around 100 ms following the deviant stimuli. This information is then used and the corresponding source is found by fitting a dipole to the interval around the peak of the difference waveform. It is noteworthy, that in the classical MMN paradigm we ideally would like this deviation to occur only within a single time interval so that the rest of the deviant stimulus response is similar to the standard response.

### 3.1 Optimization problem

We now describe an optimization problem underlying the proposed method for detecting the hidden regularities of evoked response with known temporal structure. The goal here is to find such a spatial filter **w** that the projected differential ERP *D^*(*t*) = *Ê*^*c*1^(*t*) − *Ê*^*c*2^(*t*) = **W**^*T*^ (*X*^*c*1^(*t*) − *X*^*c*2^(*t*)) satisfies the following two conditions:

1. *D^*(*t*) should have maximum deflection within the Target range, see figure 1.
2. *D^*(*t*) should have minimal possible amplitude over the rest of the response time, the flanker range, see figure 1.

**Figure 1:**
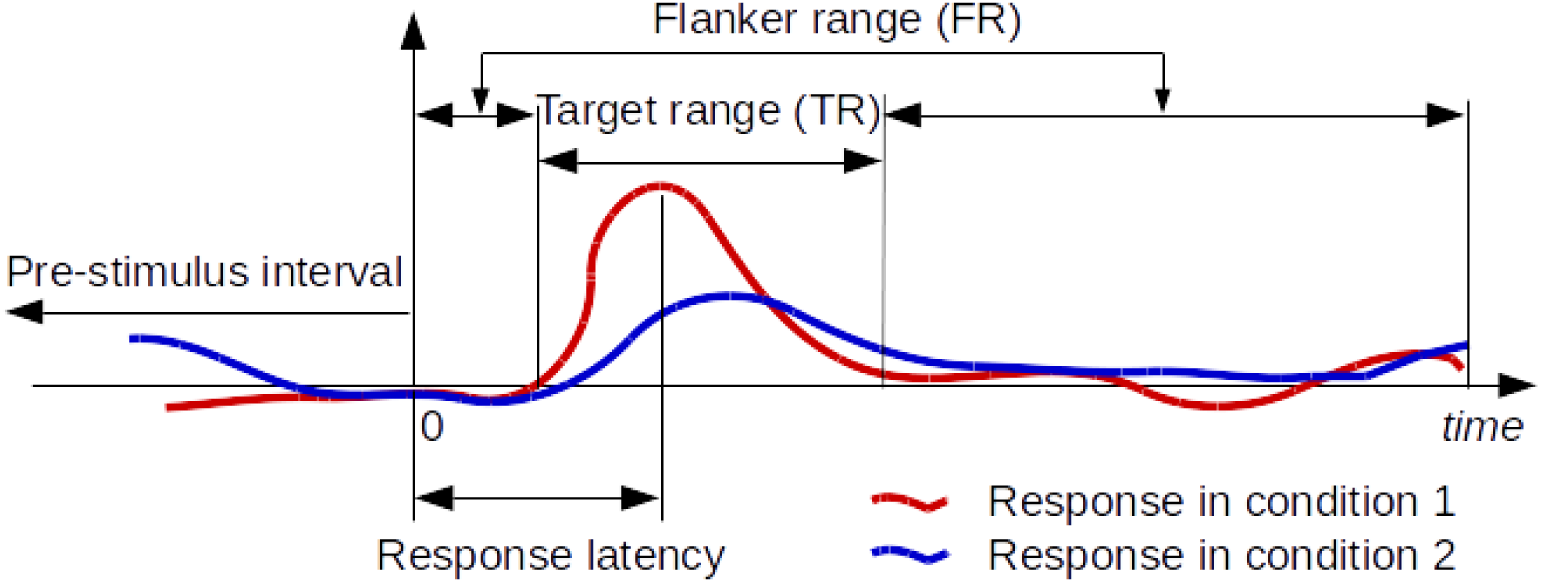
The diagram illustrating the proposed method. Two typical ERPs are superimposed. We expect to see statistically significant difference between the two conditions in the Target range and to see no such difference over the rest of the time points (flanker range)

Similarly to the CSP method, the corresponding problem can be formulated as a generalized Rayleigh quotient maximization:

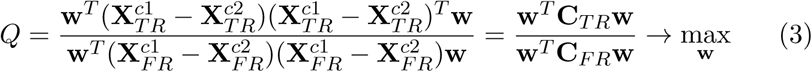

where 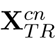 is a [*K* × *T_TR_*]-matrix containing the condition *n* ERP samples from the target range (TR), and 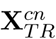 is a [*K* × *T_FR_*]-matrix containing the complimentary subset of condition *n* ERP samples that form the flanker range.

Since the intervals may contain fewer time samples than the number of channels *K* the resultant matrices **C**_*TR*_ and **C**_*FR*_ will be ill-conditioned. In order to resolve this we replace these matrices with their Tikhonov regularized versions as (**C̅**_*TR*_ = **C**_*TR*_ + λ_*TR*_**I** and (**C̅**_*FR*_ = **C**_*FR*_ + λ_*FR*_**I** where λ is taken to be a small fraction (0.1) of the *trace*(**C**)/*K* – average diagonal element. Alternatively, the shrinkage technique can be used to analytically compute a value for λ.

Thus, our method reduces to finding the **w** that maximizes the following Rayleigh quotient

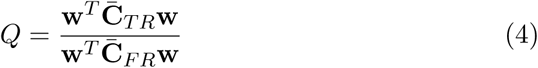

which can be performed by solving the generalized eigenvalue problem for the (**C̅**_*TR*_, **C̅**_*FR*_) pair of matrices. As a result we will obtain matrix **W** of generalized eignevectors so that (**C̅**_*TR*_**W** = **ΛC**_*FR*_**W**. The optimal spatial filter **w**\* will correspond to the largest eigenvalue of the 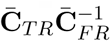 matrix. In order to find the matrix **V** of the corresponding topographies we invert the generalized eigenvectors matrix so that **V** = **W**^−1^. The topography of the source whose activity matches the desired temporal profile is then given by the corresponding row of **V**.

### 3.2 Statistical testing procedure

Since the optimal filter **w**\* computed and applied to the ERP we should provide the guarantees that the obtained component is not spurious. In order to check the consistency of obtained results the nonparametric permutation test can be used [3]. Under the null hypothesis all trials in the data are assigned with some unknown probability distribution *fX_j_* = *f*(*X_j_* = *x_j_*). The null hypothesis of the permutation test states that responses in all trials are drawn from the same distribution, regardless of the condition under which it was recorded (*c* = {*c*1, *c*2}). The alternative hypothesis states that the probability distributions are different due to the different experimental conditions.

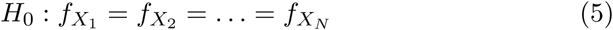

To perform the test we need to draw new samples from the permutation distribution, apply the statistical method proposed to the new samples and compute some statistics *S* which can be compared with the same statistics for the original data *S*\*. After *M* repetitions of permutation resampling, the p-value of the test computed as a ratio of the statistics values that exceed the original value

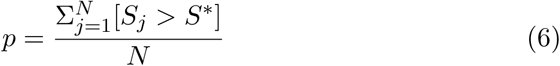

Drawing from the permutation distribution implies randomly permuting trials under the different experimental conditions in such a way that all possible permutations appears with equal chances. The statistics should reflect the difference between the responses observed in different conditions. If the value for permuted samples is higher than for the original data it means that the detected effect is spurious and does not evoked by the experimental conditions.

## 4 Data description

In this paper we describe an application of the proposed statistical method to the data acquired in the study devoted to the analysis of the auditory neuroplasticity of neuroeconomics. During this study subjects participated in two sessions of the experiment in two consecutive days. The experimental paradigm combines two widely used approaches: monetary incentive delay task (MID task) and oddball session.

The design of the study has several specific features. While in the original MID task the incentive cues are visual, the goal of this research is analysis of the auditory neuroplasticity, so the auditory modification of MID task were used. During two consecutive MID task sessions the participants built associations between sounds and monetary rewards they could get in each trial corresponding to the sound. Since the design includes two oddball sessions – one before the MID task and one after - the differences in the mismatch negativity (MMN) component magnitudes between two days can be used as a marker of neuroplasticity caused by learning.

During the oddball session the subject heard sequences of sounds, where on of them was the standard sound and appears much frequently than other deviant sounds. The sounds differs both in frequency and intensity and during the session all deviants were presented in the randomized sequence. The number of each deviant amounts to 5% of the standards. EEG data were recorded using 60 active electrodes (Brain Products GmbH) at a sampling rate of 500 Hz.

It is suggested that MMN changes its magnitude as a result of learning only in the case when subject did not discriminate the sounds absolutely before the experiment [5]. Therefore, the sounds used as incentive cues in the MID task and as deviant sounds in the oddball session were chosen in accordance with a personal auditory sensitivity level and were not absolutely discriminated by the subjects.

Unexpectedly, the observed MMN magnitude in the acquired data is too small to detect it on the single channel. It seems that the possible explanation of this phenomena is that the sounds were too close to each other and it was difficult for a subject to distinguish them.

## 5 Results

In order to detect the MMN component in the data described above we use the proposed statistical method of maximal discrimination between evoked responses in the time interval of interest. The developed method implies single-trial within-subject analysis, so here we demonstrate the results on the one subjects’ data.

As in this case we are looking for the MMN, the Target range for calculations is 80–180 ms since it covers the period when we expect to observe the component. After the filter **w**\* was calculated, the permutation test was performed to check the consistency of the results. In the described case the null hypothesis of the permutation test is rejected at the failure rate of less than 5% (p-values lower that 5%).

Figure 2 demonstrates the projected responses with filter 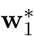 for the first day (blue line) and 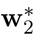 for the second day (red line) after the training. The Target time interval of 80–180 ms is highlighted. It is clearly seen that the filtration allows us to detect the significant peak in the Target interval while the difference in responses in flanker range fluctuates around 0. The topographies of the potential spatial distribution are demonstrated on the B) and D) parts of figure 3.

**Figure 2:**
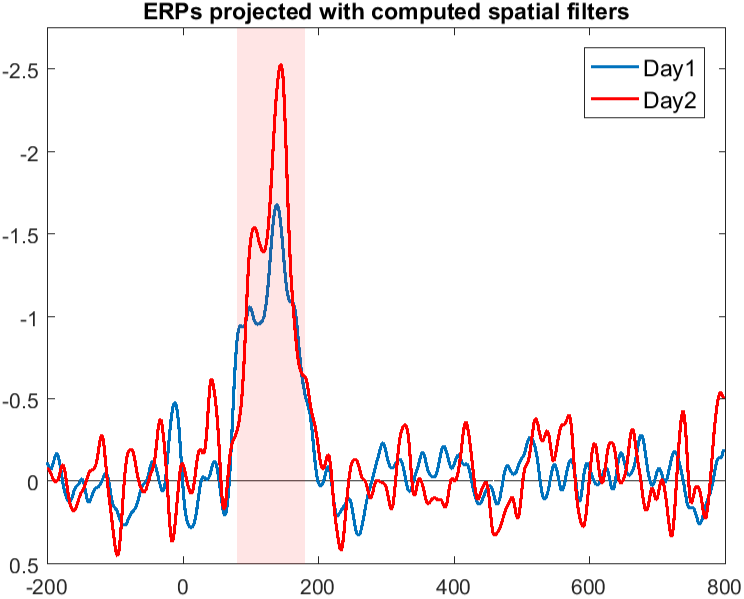
Difference between averaged deviant trials and standard trials projected with the obtained spatial filters; the highlighted time interval is 80–180 ms used as a target interval for discrimination of two conditions.

**Figure 3:**
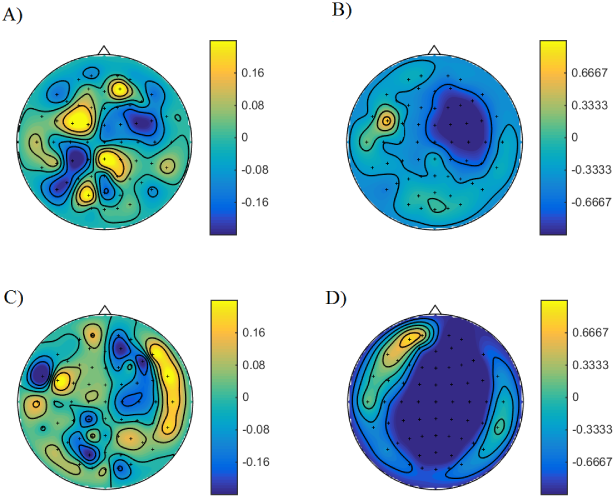
A) Spatial distribution of filters computed (Day 1); B) Topography of the potential distribution (Day 1); C) Spatial distribution of filters computed (Day 2); D) Topography of the potential distribution (Day 2);

Since the detected component has an appropriate latency and negative potential deflection is located in the fronto-central area, the detected component can be defined as MMN. We can conclude from the figures that this subject were successfully trained during the MID-task as both amplitude and area of propagation of the component increased on the second day.

Consistency of the obtained results is supported by the analysis of the whole time interval with the proposed method. We fixed the length of Target range for 100 ms and then moved the starting point from the beginning of the whole range (−200 ms) to the end (700 ms) with step of 20 ms. Figure 4 demonstrates the maximal obtained eigenvalues for each interval for two days. Figure 5 shows the corresponding p-values from the permutation test performed for chosen target time intervals, horizontal dashed lines cut the classic thresholds for p-values of 0.1, 0.05 and 0.01.

**Figure 4:**
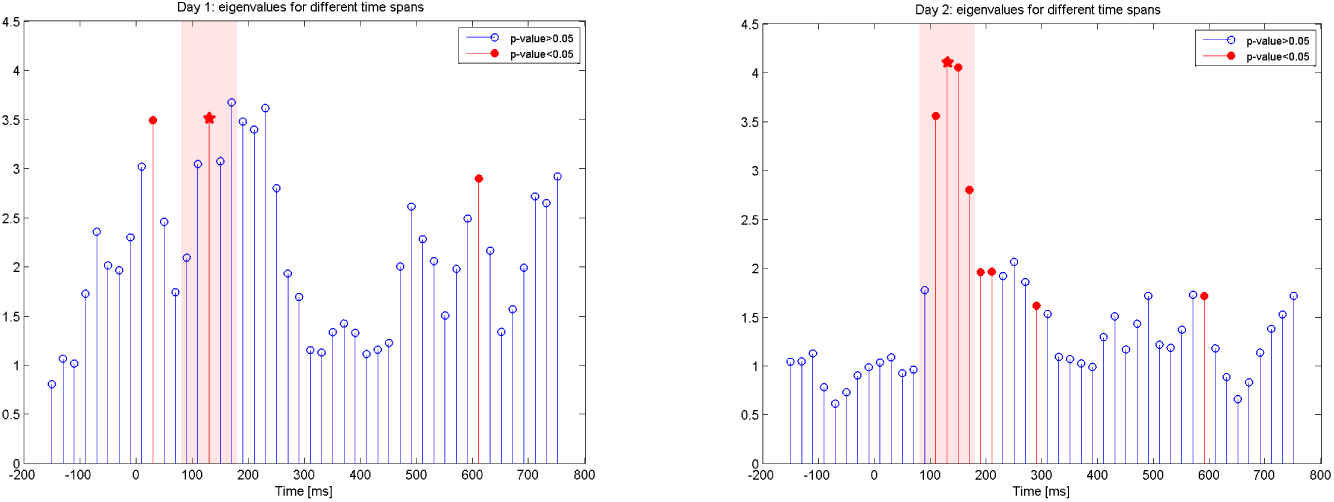
Eigenvalues for different time spans used for filter calculation, two days. The highlighted region is 80–180 ms. Eigenvalues colored in accordance with the statistical significance of the observed component (red points correspond to the permutation test p-values < 0.05).

**Figure 5:**
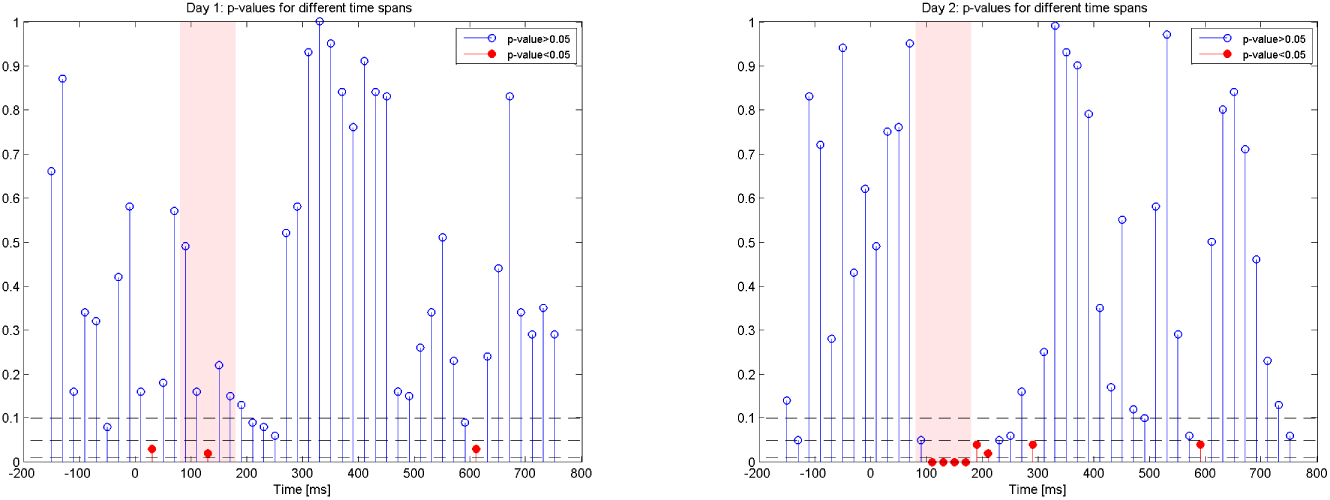
Permutation test p-values for different time spans used for filter calculation, two days. The highlighted region is 80–180 ms. Dashed lines cut off the 0.1, 0.05 and 0.001 levels, points under the 0.05 threshold colored in red.

Magnitudes of maximal eigenvalues equal to the maximal value of the Rayleigh quotient (equation (4)) obtained with the optimal filters **w**\*. The correct Target range which used for MMN detection previously highlighted with red and the eigenvalues corresponding to this time span marked with stars. While there are a lot of eigenvalues which are rather high in comparison with the target eigenvalue (in the true Target value 80–180 ms), the permutation test approve the consistency only for three of them (colored in red).

The evidence of the MMN magnitude increase on the second day are supported by the fact that there are several time intervals concentrated near the highlighted initial Target range, which give the significant result after the filtration. This effect can be explained by the enlarged propagation of the MMN component in time.

## 6 Acknowledgements

The study was supported by the grant 16-18-00065 of the Russian Science Foundation.

